# In situ characterization of late Chikungunya virus replication organelle

**DOI:** 10.1101/2023.02.25.530016

**Authors:** Justine Girard, Olivier Le-Bihan, Joséphine Lai-Kee-Him, Maria Girleanu, Eric Bernard, Cedric Castellarin, Aymeric Neyret, Danièle Spehner, Xavier Holy, Anne-Laure Favier, Laurence Briant, Patrick Bron

**Affiliations:** CBS (Centre de Biologie Structurale), Université de Montpellier, CNRS, INSERM, Montpellier, France; IRIM (Institut de Recherche en Infectiologie de Montpellier), Université de Montpellier, CNRS, Montpellier, France; IRBA (Institut de Recherche Biomédicale des Armées), Ministère de la Défense, Brétigny-sur-Orge, France

**Keywords:** Alphavirus, Chikungunya, Replication organelles, nsP1, cryogenic-electron microscopy, sub-tomogram averaging

## Abstract

Chikungunya virus (CHIKV) is a mosquito-borne pathogen responsible for an acute musculo-skeletal disease in humans. The viral RNA genome replication occurs in membrane spherules named replication organelles (ROs). In this work, we investigate the native structural organization of CHIKV ROs in their cellular context using *in situ* cryogenic-electron microscopy approaches at late replication stage. We observed previously unreported diameter heterogeneity of ROs at the plasma membrane of infected human cells. CHIKV ROs were only marginally detected in cytopathic vacuoles where they are homogeneous in size, suggesting a finely regulated internalization process. Our data show that ROs maintained at the plasma membrane beyond the first viral cycle are dynamically active both in viral RNA replication, in its export to the cell cytosol, but also in the production of viral proteins. We suggest that late CHIKV ROs have an amplifying role or represent an alternative pathway in the production of infectious viral particles. All these observations bring new insight into the CHIKV life cycle in human cells.

## INTRODUCTION

Positive-stranded RNA viruses replicating in the host cytoplasm dramatically remodel intracellular membranes into specialized vesicular structures supporting viral RNA genome replication and transcription ^1,21,52^. Generally separated from the cytoplasm by a proteinaceous pore, these viral replication organelles (RO) provide an optimal micro-environment that concentrate metabolites, and viral and host components required for genome replication. They also presumably shield double-stranded RNA (dsRNA) replication intermediates from innate immune sensors and antiviral effectors, and may additionally coordinate genome replication, viral translation, and new particle assembly. Deciphering ROs organization, biogenesis, and mechanisms of maintenance at the molecular level, therefore, represents an area of intense interest with potential consequences for therapeutic intervention. The recent advance in cryogenic-electron microscopy (cryo-EM) and cryogenic-electron tomography (cryo-ET), sub-tomogram averaging and 3D reconstruction techniques has provided a substantial advance in elucidating the 3D volume architecture of ROs assembled by a variety of pathogenic RNA viruses (e.g. nodaviruses, flaviviruses, bromoviruses, tombusviruses, coronaviruses) ^2–9^. Contrasting with this significant breakthrough, our knowledge of *Alphavirus* ROs is still incomplete.

*Alphaviruses* (Togaviridae family) are mosquito-borne viruses that can cause a severe human illness including persistent arthritis and fatal encephalitis. Among these, the Chikungunya virus (CHIKV), responsible for severely debilitating and often chronic rheumatic disease ^10–12^, has become a major public health issue, notably due to the rapid spreading of its mosquito vectors *Aedes aegypti* and *Aedes albopictus* ^13,14^. CHIKV is an enveloped virus of about 50-60 nm in diameter ^15^. Its replication cycle has been the focus of an intense attention, resulting in the following picture ^16–18^. Replication is initiated when the viral genome, a positive-sense single-stranded RNA molecule of 11.8kb containing a 5’-cap and a 3’ polyadenylated tail is loaded by the host translation machinery. The 5’-proximal two-third of the viral genome corresponding to the first open reading frame (ORF) is translated into a nonstructural polyprotein precursor (P1234) which associates with an RNA genome and is trafficked to the plasma membrane ^19^. P1234 sequential proteolytic processing results in the release of the four nonstructural proteins (nsP1 to 4), which assemble to form the viral replicase. During this reaction, this complex replicates the RNA genome through the synthesis of a negative-strand RNA template ((-)RNA), resulting in the accumulation of double-stranded RNA (dsRNA) replication intermediate species ^20,22^. Starting from the (-)RNA, CHIKV replicase also controls the transcription of a 26S subgenomic RNA (sgRNA) corresponding to the 3’ ORF, translated into five structural proteins: capsid (C), envelope and surface glycoproteins (E1, E2, E3), and 6K/TF, required to form the nascent viral particle.

Alphavirus genome replication takes place in bulb-shaped ROs, referred to as spherules first assembled at the plasma membrane (PM) ^1,21,23,24^. For some alphaviruses, these compartments undergo endocytosis to form intracytoplasmic vacuolar structures, positive for lysosomal and endosomal markers, referred to as cytopathic vacuoles (CPV) ^25^. Membrane targeting of the entire replication complex (RC), composed of nsP2, the RNA helicase and cysteine protease responsible in polyprotein processing ^26,27^, nsP3, which contains an ADP-ribosyl binding and hydrolase activity ^28^ and associates with a variety of proviral cell factors ^29^, and nsP4, the RNA-dependent RNA polymerase ^30^, is guided by nsP1, the viral capping enzyme_ ^21,31^. This 540-long amino acid methyl/guanylyltransferase behaves as a monotopic protein with a preference for anionic lipids and cholesterol-rich microdomains ^32–34^. Mutagenesis investigations assigned such affinity to the α-helix fold of the central part of nsP1 and to the presence of palmitoylated cysteines which, despite not being required for membrane anchoring, target nsP1 to lipid rafts and support nsP1’s capacity to reshape the cell membranes ^32–35^. Membrane binding is critical for activation of nsP1 S-adenosyl-L-methionine (SAM)-dependent methyltransferase (MTase) and m^7^GTP transferase (GTase) activities which ensure viral RNA 5’ cap synthesis ^34,36,37^. Recent ultrastructure analysis provided critical insights in the organization of recombinant nsP1 interacting with artificial membranes ^38,39^. These studies revealed nsP1 capacity to assemble into dodecameric rings of 14 nm in diameter with a central pore of 7.5 nm, interacting with the lipid bilayer. This proteinaceous ring was proposed to contribute to the structural maintenance of replication compartments and to mediate export of newly synthesized RNA to the cytosol where they are translated or trafficked to assembly sites. The location of the active enzymatic site to the internal face of this ring is finally thought to ensure the simultaneous capping of the multiple RNA molecules trafficked through this pore. Despite this recent knowledge, the validity of this model in infected cells as well as the global *in situ* spherule organization remains unclear.

In the present work, we have investigated CHIKV spherules structural organization in their cellular context using *in situ* cryogenic-electron microscopy approaches, combined with sub-tomogram averaging approach. To access a dynamic view of CHIKV replication cycle, we imaged the infected cells at 17h post-infection (hpi), a time point that encompasses a second and possibly a third CHIKV replication cycle that is 6-8h long. We observed CHIKV ROs as single membrane vesicles in continuity with the plasma membrane, present both on the cell body and filopodia-like extensions. Contrasting with previous electron microscopy observations performed after one viral cycle (6-8h), we observed a wide-diameter distribution of ROs. Associated with these structures, we visualized the presence of extruding filaments resembling viral RNA, tightly associated with ribosome-like densities and additional complexes nearby, arguing for a strong CHIKV translating activity at the plasma membrane. In addition, we also observed that a part of ROs, with a smaller and more homogeneous diameter, is internalized into cytosolic vacuoles. The sub-tomographic averaging of the RO neck evidenced that, in infected human cells, nsP1 forms a ring compatible with that assembled *in vitro* ^38,39^ strongly interacting with the PM lipid bilayer and provides additional details for CHIKV ROs organization in human cells.

## RESULTS

### CHIKV replication sites preferentially localize at the plasma membrane of human epithelial cells

*Alphaviruses* ROs, which concentrate the four virus-encoded nsPs and double-stranded RNA (dsRNA) replication forms are first assembled at the plasma membrane (PM) of the infected host cell, and later internalized and fused with endosomal compartments to produce type-1 cytopathic vacuoles ^25^. We first questioned the localization of CHIKV replication sites in the virus permissive HEK293T epithelial cell line infected with a CHIKV reporter virus (BNI-CHIKV_899 strain) containing a mCherry coding sequence in nsP3 hypervariable domain ^40^. To increase the probability to detect replication events, we used an MOI of 50, and to access the different replication steps the cells were infected for 17h, a duration that allows several replication cycles (estimated for alphavirus to around 6-8h) to occur ^11,41^ (Figure S1). Replication sites were identified by immunostaining of the four nsPs together with dsRNA, which represents a good RCs marker, and by confocal microscopy. In our experimental conditions, colocalized fluorescence signal is mainly detected at the plasma membrane (PM) (Figures S1A and S1B). Interestingly this signal is observed in cellular extensions detected in these cells (Figure S1B, merge), suggesting that replication compartments take place both on the cell body and at the PM limiting virus-induced filopodia-like extensions previously reported in *alphavirus*-infected vertebrate cells ^42^. Finally, a minor fraction of dsRNA colocalizes with both nsP1 and nsP3 in cytosolic aggregates, supporting that CHIKV replication complexes undergo some marginal endocytosis events as suggested ^43^. According to these characteristics, CHIKV replication sites detected at the plasma membrane of cell bodies or cellular extensions are suitable for cryo-EM investigation.

### Cryo-electron tomography of CHIKV ROs reveals their colocalization on cell body and filopodia-like extensions plasma membrane of infected cells

HEK293T cells were directly grown onto gold EM grids, infected with CHIKV, and embedded in vitreous ice. Compared to confocal experiments, we increased cell density to maintain sufficient cells onto grids after their freezing, still keeping a high multiplicity of infection (MOI 50) and 17h of CHIKV infection. We collected more than 300 cryo-tomographic tilt series of the infected cell peripheries that were processed in etomo ^44^ and segmented using Amira software (Thermo Fisher scientific) (Figure 1). These tomograms of CHIKV-infected cells are in agreement with confocal observations, revealing the presence of numerous filopodia extending from HEK293T cell bodies. The ROs appear as clusters of round vesicles of variable diameter covering the PM. They are delimited by a lipid bilayer connected to the PM through a narrow neck (Figure3 1A and 1C). Analyzing spherules content, all compartments imaged contain a compactly coiled filamentous density which corresponds to dsRNA packed inside as previously observed for Nodavirus ROs ^3^. No additional inner density is observed, indicating the absence of an internal coat contributing to membrane curvature. Contrasting with this appearance, CHIKV virions are observed as spherical particles of about 50-60 nm in diameter, highly electron-dense, and covered by protruding densities (Figure 1C). In the present study, most CHIKV replication spherules are located at the PM limiting the cell body of infected cells (Figure 1A) while filopodia-like extensions exhibited some ROs colocalized with most CHIKV budding particles (Figure 1C). The computed 3D reconstructions granted us access to the connection of ROs with the PM and other cellular structural details of ROs local cellular environment, notably revealing the presence of cytoskeleton filaments or cell host factors like ribosomes. When both ice thickness and defocus values used during the tilt series acquisitions are low, the level of details observed in images can be high. Thus, it is possible to discriminate the double-layer organization of the plasma membrane or observe the repetitive nature of actin filaments present in filopodia-like extensions (Figure 1C), some good indicators of the datasets intrinsic resolution. It is worth noting that CHIKV infection induces local cellular re-arrangement. While filopodia normally appear as cellular extensions containing a continuous bundle of parallel actin filaments in close contact with the PM, actin filaments in CHIKV-induced filopodia-like extensions bearing ROs and/or CHIKV viral particles are separated from the PM by a space in which cellular compounds like ribosomes accumulate (Figures 1 and 2 versus S2). This phase separation suggests a cellular activity that certainly relies on the CHIKV viral cycle, possibly associated to the production and assembly of viral particles. Indeed, as exposed in Figure 2 (blue arrows), filopodia-like extensions also display CHIKV budding sites near ROs, often associated with a thickening of the PM, a characteristic recently correlated to the insertion of immature envelope spikes in the lipid bilayer ^45^ (Figure 2, yellow arrows). Furthermore, ROs from the aforementioned region bear some extra densities protruding out of their membrane (Figure 2, red arrows). Considering their proximity to budding sites, these densities likely correspond to CHIKV envelope glycoproteins. This unexpected feature is in agreement with recent immuno-labeling experiments of CHIKV-infected cells with anti-E2 antibodies reported by Jin et al. ^46^, revealing some signals at the surface of ROs.

**Figure 1:**
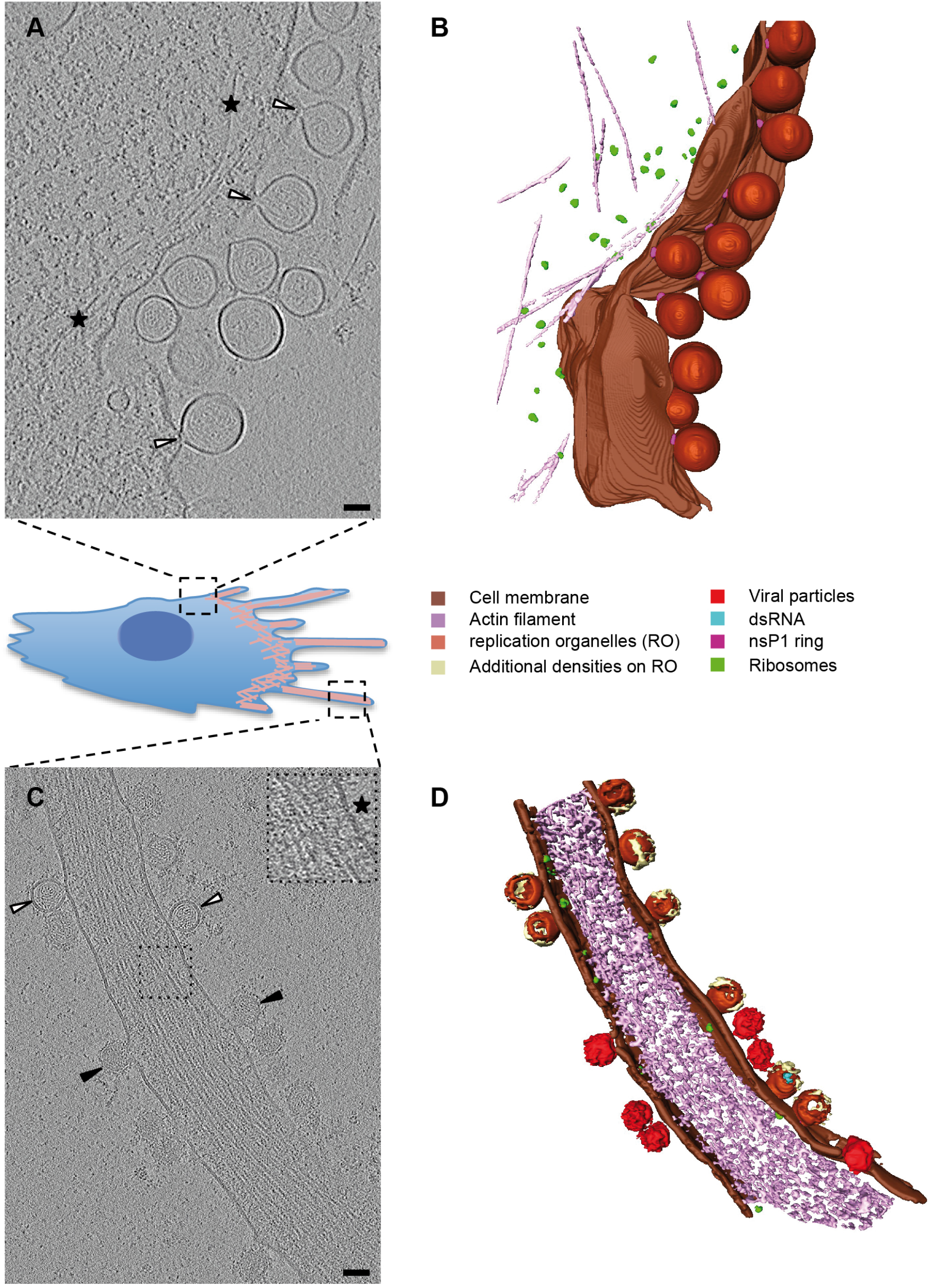
The CHIKV replication organelles are observed at the plasma membrane of limiting cell body and filopodia-like extensions of infected cells. **A and C.** Sections in electron tomogram of a HEK293T cell infected for 17h at MOI 50. Spherules are connected to the PM through a narrow neck (white arrows). Some cytoskeleton filaments present beneath the PM are indicated by black stars. **B and D.** Segmentation of (A) and (C) showing the presence of CHIKV ROs (white arrows) and budding CHIKV viral particle (black arrows) onto the same cell filopodia-like extension. The bilayer organization of the PM (black star) and the repetitive nature of actin filaments (low-pass filtered inset) are clearly visible. Scale bar; 50 nm.

**Figure 2:**
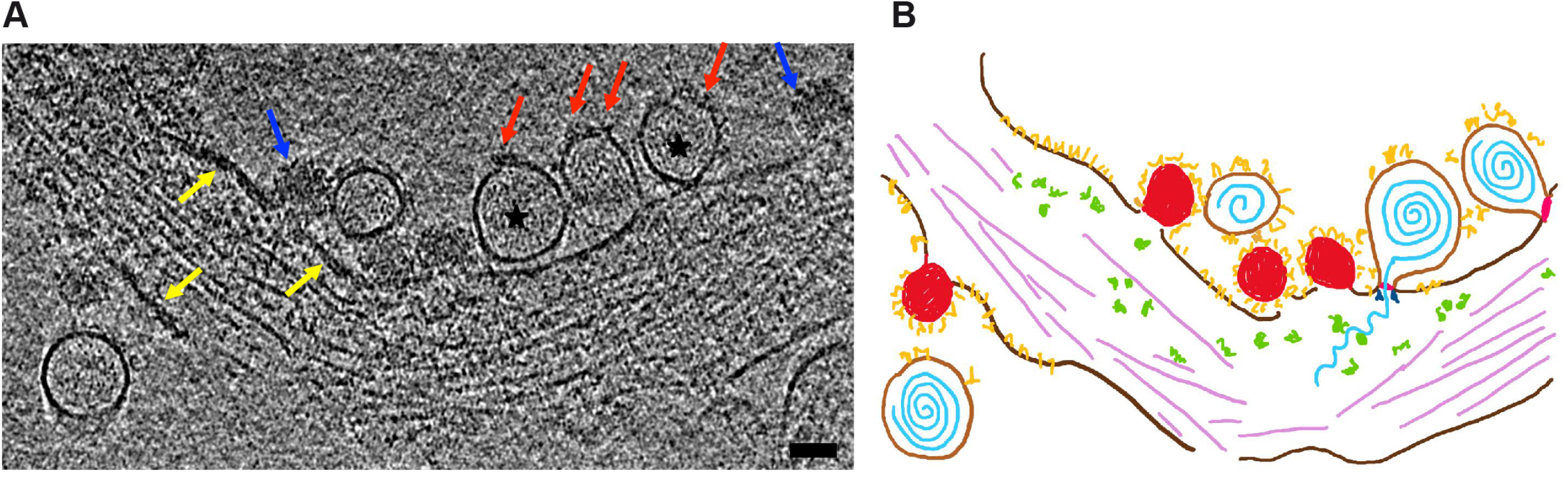
Tomogram section of CHIKV-induced filopodia-like extension. **A.** Electron tomogram of a CHIKV-induced filopodia-like extension. While CHIKV particle are detected at the PM (blue arrows), CHIKV ROs appear less dense and larger (black stars). Some additional densities are anchored into ROs limiting membrane (red arrows). A thickening of the PM can appear locally (yellow arrows). Scale bar, 50 nm. **B.** Schematic representation of A. Actin filaments are represented in light purple, ribosomes in green, cell PM in brown, viral particles in red, CHIKV ROs in light brown, the dsRNA in light blue, immature spikes in yellow, nsP1 putative ring in pink, and some replication partners in dark blue.

**Figure 3:**
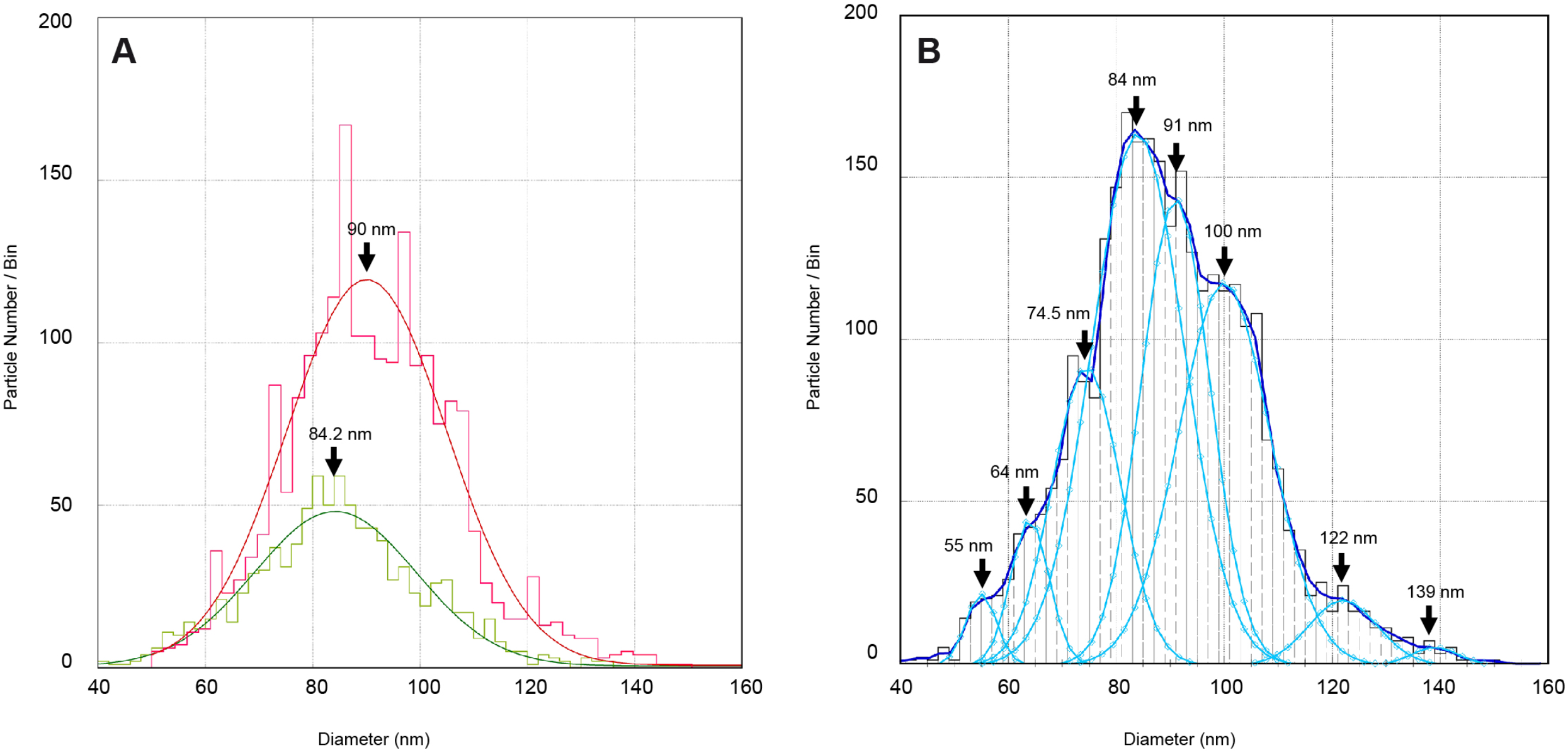
CHIKV replication organelles diameter distribution. **A.** Histogram distribution of ROs diameters computed from 2,083 measurements for ROs located on cell bodies (red) and 950 measurements for ROs located on cell filopodia-like extensions (green). Both populations fit with a Gaussian distribution with a mean diameter of 84.2 and 90 nm for cell body and filopodia-like extension ROs respectively. **B.** Histogram distribution of ROs diameters considering all ROs, corresponding to 3033 measurements. Curve fitting deconvolution unveils eight Gaussian sub-populations whose mean diameters are indicated.

### CHIK ROs display variable size distribution

Alphavirus ROs have been initially reported to display a standard diameter of around 50 nm ^1,47^. As mentioned above, CHIKV replication compartments detected in our experiments primarily appeared unexpectedly heterogeneous in size (Figure 1), an observation also validated by CHIKV infected cells plastic sections performed at 17 hpi at 4 MOI (Figure S3). To objectify this observation, we measured the diameter of more than 3000 ROs from cryo-tomograms. As ROs can be elongated perpendicularly to the PM, we considered the maximal diameter of ROs parallel to the PM (Figure S4). In this context, we determined the diameters of 2,083 and 950 ROs from PM of cell bodies and filopodia-like extensions respectively, and plotted them as a histogram representation (Figure 3). RO diameters display a Gaussian distribution ranging from 40 to 140 nm with a mean of 84.2 nm and 90 nm for ROs located at the PM of cell bodies and filopodia-like extensions respectively (Figure 3A). Thus, we only observed a slight shift of the mean diameter between ROs localized on the PM of cell bodies and filopodia. It is worth noting that we never observed ROs with a diameter small than 40 nm. Moreover, the biggest ROs, having a diameter superior to 120 nm, are only found at the PM of cell bodies. While these observations support that ROs assembled at the PM of cell bodies can be slightly bigger than those detected on filopodia, this feature is not entirely clear, especially due to the overall unexpected ROs size variability observed in our experiments.

In a closer analysis of our data, considering the whole ROs population, we discerned several RO diameter maxima of 55, 64, 74.5, 84, 91, 100, 122, and 139 nm (Figure 3B). It is worth noting that some very recent cryo-EM studies reported RO with diameters of about 50 to 70 nm formed 6-8h after CHIKV infection ^48,49^, a time point at which CHIKV first replication cycle is admitted to be complete, resulting in the presence of mature ROs containing fully-processed nsPs and dsRNA replication intermediates. Contrasting with this experimental scheme, our observations were set at 17h after the CHIKV challenge. Previous molecular studies established that the volume of ROs and consequently their diameter tightly correlates with RNA genome length and indirectly with the size and amount of neosynthesized genetic material contained in these compartments ^3,50^. Considering this information and to the late time point considered in this study, we propose that ROs having a diameter superior to 80 nm may correspond to ROs in which the viral RNA continues accumulating. Accordingly, the idea of finite final size of ROs needs to be reconsidered according to the post-infection time.

### CHIKV internalization in cytopathic vacuoles is regulated

Our confocal microscopy and electron microscopy analysis reveal that after a prolonged CHIKV infection, most ROs remain at the PM. To confirm this peculiarity and investigate the presence and structural organization of ROs internalized in CPVs, CHIKV-infected cells were processed according to the cryo-EM method of vitreous sectioning (CEMOVIS) ^51^. CPVs were rarely detected in vitreous sections, and usually at vicinity of mitochondria (Figure 4). They appear as vacuolar structures ranging from 200 to 500 nm in size, delineated by a lipid bilayer and containing budded spherules. From a morphological point of view, these organelles are similar to ROs located at the PM with a lipid membrane delimiting an internal area full of ball-of-yarn-like densities and connected to the CPV membrane envelope via a neck. No obvious additional density could be observed in the cytoplasm beneath these spherules. Although we did not observed in our vitreous sections, CPVs are often associated with honeycomb arrangement of capsid proteins (Figure S5) (José et al. 2017). Contrasting with previous observations on related Alphavirus ^1,41,52^, CHIKV spherules appear disorganized in these compartments, instead of lining the vacuole membrane, a feature that is also observed in plastic sections observed by conventional electron microscopy (Figure S5). Interestingly, in these compartments, ROs display a rather homogeneous diameter of about 50.6 nm ± 6 nm (Figure 4), therefore contrasting with the heterogeneous size detected at the PM. This unexpected homogeneity in size argues for a regulated internalization mechanism susceptible to occur at a defined step of CHIKV replication cycle.

**Figure 4:**
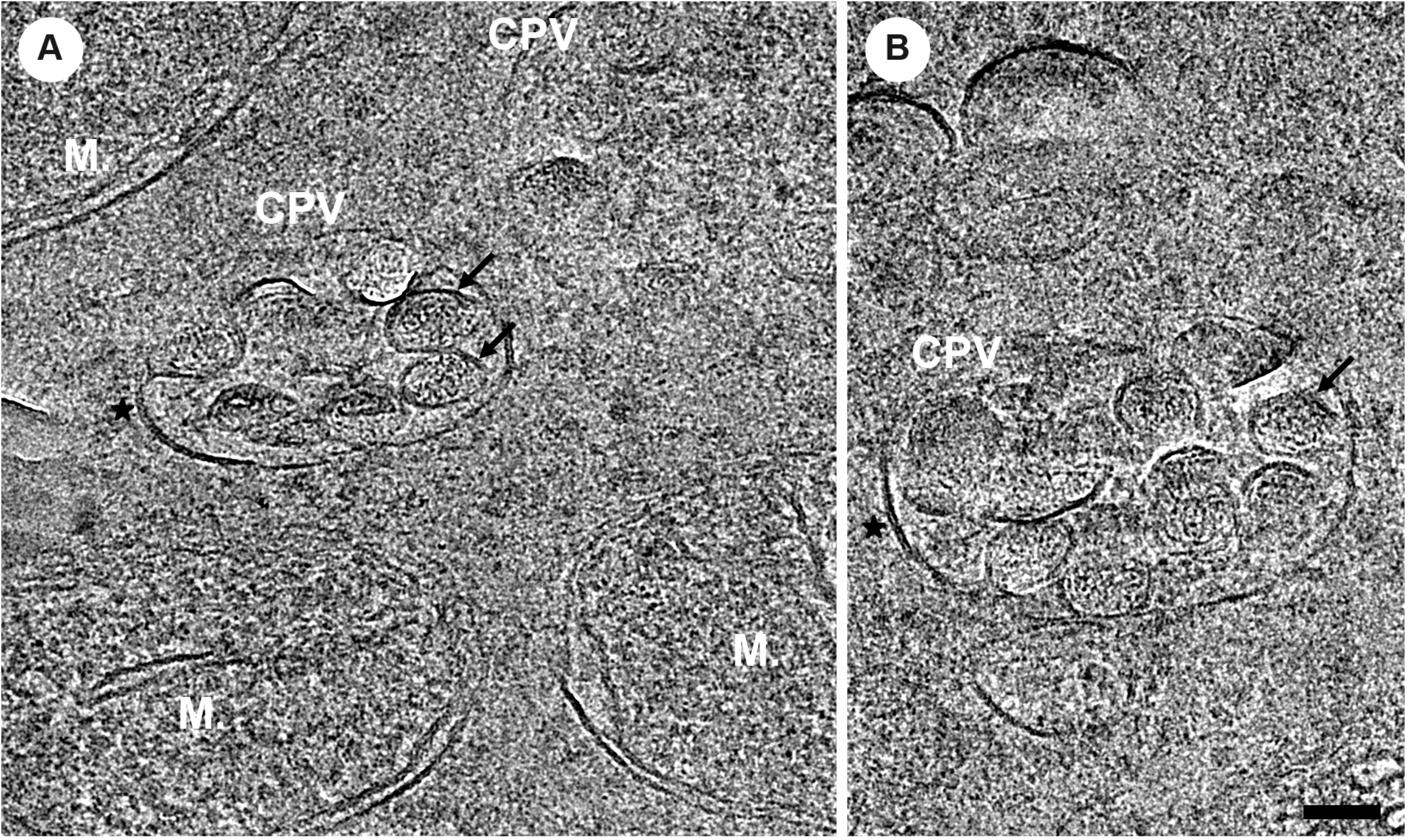
Cryo-EM vitreous sections of CHIKV infected cells reveal the presence of cytopathic vacuoles. **A and B**. Some RO-filled cytopathic vacuoles (CPV) (black arrows), 200 to 500 nm in diameter are observed in vitreous sections. They are delineated by a lipid bilayer and often found at vicinity of mitochondria (M). Scale bar, 50 nm.

### Late CHIK ROs exhibit heterogeneous patterns

Considering the above-discussed ROs size heterogeneity, the functionality of compartments with an unexpected diameter may be questioned. In this context, we performed a thorough inspection of our cryo-tomograms intending to report all different types of molecular patterns of ROs, taking into account the size of these organelles and the presence or absence of densities inside and immediately below spherules. In Figure 5, we display different views of ROs according to their diameter, which ranges from 50 to more than 120 nm. This exploration reveal that whatever the size of ROs, the lumen of spherules is full of ball-of-yarn-like densities, with a state of compaction that seems similar, suggesting a high RNA replication activity inside these compartments. It is worth noting that all ROs display a clear density at the base of spherules which separates the inner space of spherules from the cell cytoplasm and that seems in continuity with the PM. Thus, spherules seem to stick on the PM. Exploration of cytoplasmic areas immediately beneath ROs revealed variable patterns, varying from the absence of clear associated densities below spherules (Fig 5A; thumbnails 1 to 5) to the presence of compact or filamentous densities extruding from ROs (Fig 5B; thumbnails 6 to 12). Therefore, elongated densities can be observed just beneath spherules, displaying various length ranging from 10 nm (Fig 5, thumbnail 6) to more than 100 nm (Fig 5, thumbnail 8-10, 12). Some of these filamentous objects are covered by small globular densities, forming a sort of pearl necklace (Thumbnail 8). This pattern reminiscent of RNA covered with translating ribosomes suggests a local translating activity ^53^, which is in agreement with the local presence of CHIKV budding particles and immature envelope spikes accumulation in close proximity as in Figure 2. It is important to note that these long exported viral RNA molecules were only observed for ROs larger than 90 nm in diameter. Just beneath some spherules, two elongated, short and parallel densities can be observed, perpendicular to the PM plane and located on either side of the neck as in thumbnails 7 to 9 (in dark blue in the schematic representations). Finally, additional isolated globular densities are observed, possibly attesting to the presence of translated viral proteins and/or cell host factors recruited to replication sites. Interestingly, no clear correlation could be made between these various ROs profiles and compartment size. Altogether, these different patterns of ROs deviating from the canonical 50-70 nm size, strongly suggest that ROs maintained at the PM after the first viral cycle display viral RNA replication, its export and translating activities.

**Figure 5:**
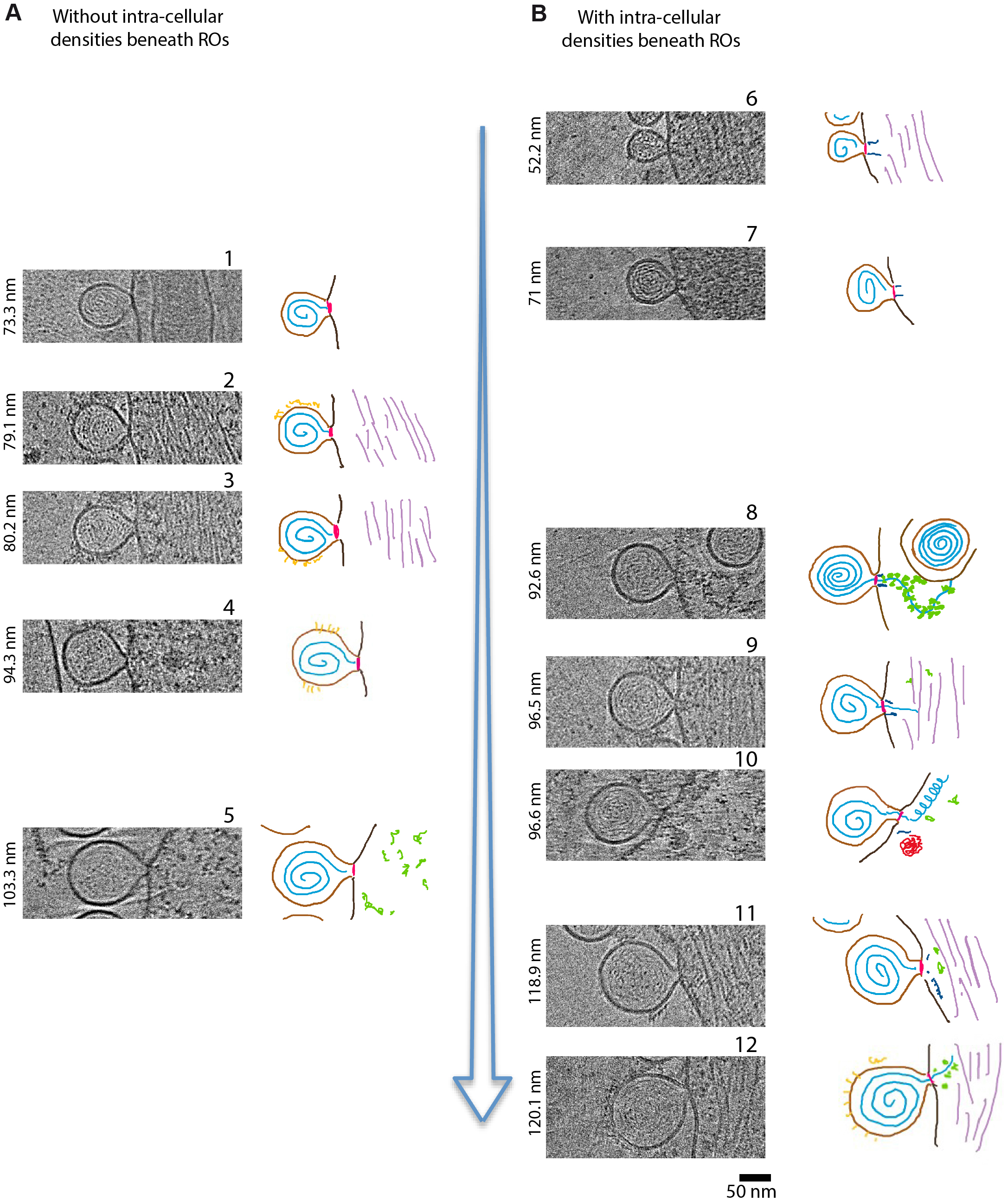
CHIK ROs display numerous patterns. ROs can be sorted according to their diameter, ranging from 40 to 140 nm and to the absence (A) or presence (B) of clear associated density patterns, just beneath the PM. A schematic representation of each pattern is also presented. The PM is drawn in black, the spherule membrane in brown, the envelope spikes decorating ROs membrane in yellow, the base of ROs neck in red, the viral RNA in blue, ribosomes in green, intracellular filaments in purple and viral or cell host partners in deep blue.

### The CHIK ROs display a crown-like structural organization of the spherule neck

ROs assembled by (+)RNA viruses are though to be connected to the cytoplasm by a pore complex localized at the spherule neck, allowing the import of metabolites and cell factors, and facilitating the export of newly synthesized viral RNA to the cytoplasm for translation and packaging. Two recent cryo-EM single particle analysis studies revealed that recombinant alphavirus nsP1 assembles into a dodecameric ring that is proposed to take place at the spherule neck ^38,39^. We investigated the possibility of transposing this model to our infected cultures at late replication stages. As previously mentioned above, we noticed the presence of a strong density at the base of each spherule neck, sometimes giving the impression that the PM of the infected cell is continuous and that ROs are just sitting on top of it. To decipher the molecular organization of the connecting region, we developed a sub-tomogram averaging (STA) approach focused on ROs neck. The STA workflow is described in details in Figure S6. Briefly, a total of 463 sub-volumes were extracted and subjected to iterative alignment, averaging and classification steps, using full rotational symmetry, resulting in a sub-tomogram average with an estimated resolution of 25 Å (gold standard FSC, FSC=0.143 threshold) from a final subset of 98 spherules. The dedicated use of masking and classification was instrumental in sorting out spherule necks morphological heterogeneity, such as variations in spherules diameter, neck length, curvature of the membrane at the base of the neck, as well as perpendicularity of spherules main axis to the PM plane. The bilayer organization of the membrane is clearly revealed in a central slice through the final reconstruction (Figure 6A) with two linear densities corresponding to the hydrophilic lipid polar heads encompassing a dark region corresponding to the hydrophobic core of the plasma membrane. The continuity of the lipid membrane bearing two clearly defined leaflets indicate that our STA strategy was able to successfully sort out morphological heterogeneity. The reconstruction reveals a high negative curvature of the PM at the base of the neck, with an average angle of 65 degrees. Additional protein densities are tightly associated with the lipid bilayer at the neck base and extend to the cytoplasmic side. This neck complex is composed of three regions: a membrane-bound ring, a barrel-like density composed of three stacked rings present in the cytoplasmic moiety and a central elongated density passing throughout all the rings. The first membrane-bound ring has an outer and inner diameter of 18 nm and 8 nm respectively and is formed by a density showing a tilted “C-like” shape, where the opening of the “C” points toward the PM. Recently, dodecameric cryo-EM structures assembled from the recombinant nsP1 have been resolved and proposed to form a connecting pore separating the inner part of the spherule from the cytoplasm ^38,39,49^. The atomic structure of this nsP1 ring perfectly fits into the density of the membrane-bound ring evidenced in our reconstruction (Figures 6B-D). The conical shape of the ring exhibiting a larger, flared base is related to the tilted arrangement of nsP1 in the dodecamer. It is worth noting that while the PM outer layer has a continuous organization, the inner layer engages extended contact with nsP1. This is in agreement with the role depicted for MBO loop 1 and 2, corresponding to nsP1 amino-acids 200-238 and 405-430 respectively in the recombinant nsP1 ring, that forms amphipathic membrane-binding spikes penetrating by about 10 Å into detergent FC12 micelles. This tight interaction is proposed to be reinforced by a triad of palmitoylated cysteines in MBO loop 2 ^39^. Nevertheless, our observations support additional interactions of nsP1 residues on the outermost domains of the nsP1 ring with PM phospholipids. Going throughout the nsP1 ring, a second central and elongated complex displays two prominent densities, one located on the spherule side and the other more pronounced located in the cytoplasmic part, is observed. As we applied a full rotational symmetry, we cannot accurately determine the shape and the stoichiometry of individual compounds involved in this elongated density. Finally, surrounding the cytoplasmic part of this central density, the additional barrel-like assembly composed of three stacked rings does not show obvious direct interaction with the nsP1 ring or the central elongated complex.

**Figure 6:**
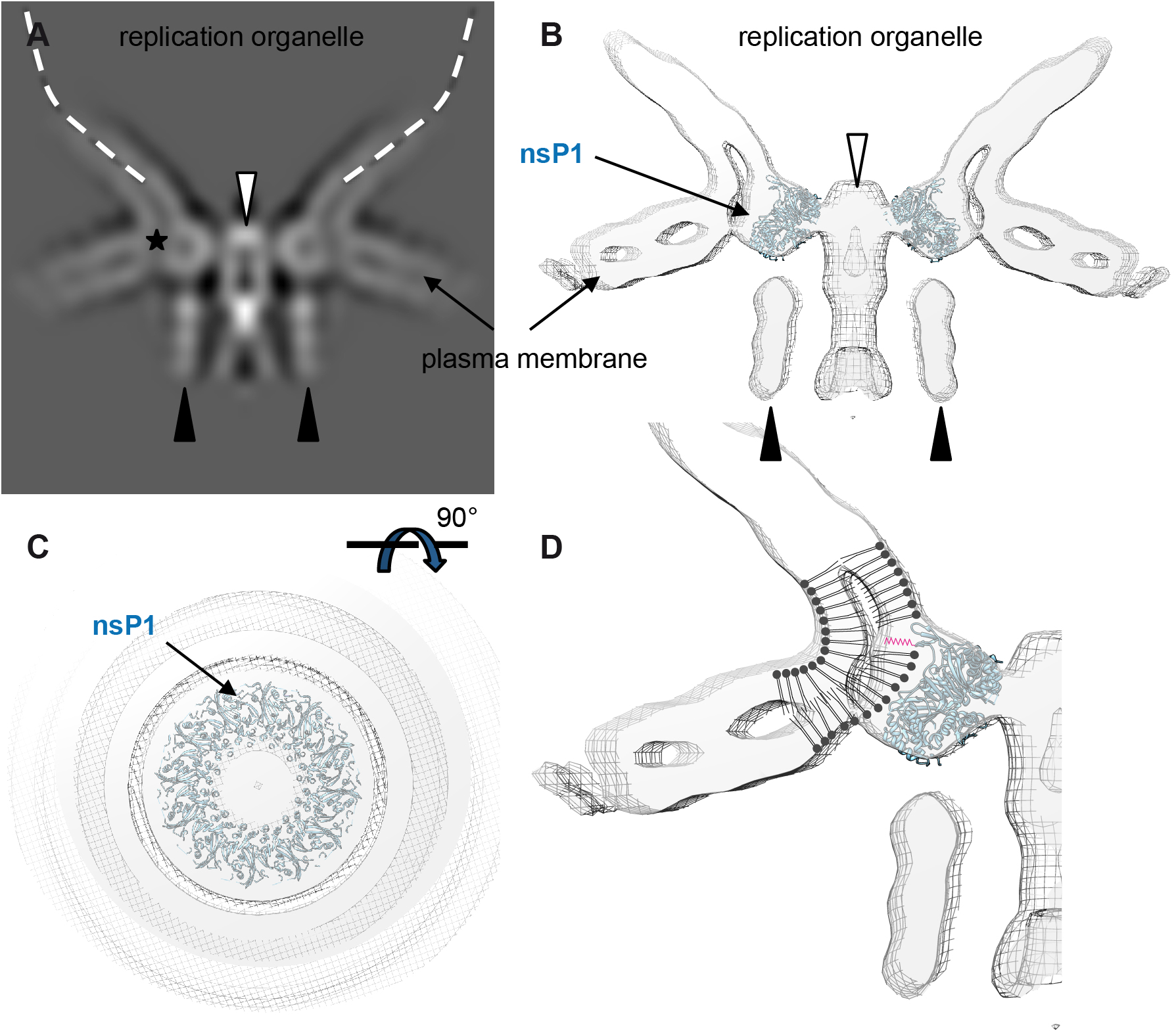
Structural organization of CHIKV RO connection to the plasma membrane revealed by sub-tomogram averaging. **A.** Central section of the 3D reconstruction. As a visual aid, the location of the spherule membrane is drawn as a white dotted line. The PM appears as two parallel white linear densities encompassing a dark region. Several densities highlighted by the white and black arrows are resolved at the level of the RO neck, as well as a C-shaped density (star) corresponding to the nsP1 ring snuggly interacting with the PM at the base of the neck. **B.** Central section of the surface representation of the RO connection to the PM. The dodecameric nsP1 ring atomic model (PDB: 6ZOV) fits the C-shaped ring density. The central channel is plugged by an elongated central density protruding toward the cell cytoplasm (white arrow) while a second tall barrel-like density composed of three stacked ring (black arrows) show no apparent connection with others densities. **C.** Surface representation of the map viewed from the interior of the spherule. A ring is present at the base of the spherule in which the dodecameric nsP1 atomic model fits perfectly. **D.** Model of nsP1 ring insertion in the PM internal layer that supports a tight interaction of nsP1 MBO loops with the PM. The insertion of the palmitoyl moities into the PM is indicated in red.

## Discussion

In the present study, with the aim at providing a new insight into the CHIKV life cycle in human cells, we investigated the native structural organization of CHIKV ROs in their cellular context using *in situ* cryogenic-electron microscopy approaches. This study considered late infection time to access the dynamic of ROs through the study of multiple infectious cycles in a single cell. Microscopy analysis of cells infected with Sindbis or Semliki Forest alphaviruses previously revealed that after initial assembly at the PM ^21^, ROs initially visualized as bulb-shaped organelles in continuity with the PM are generally rapidly internalized and fused with endosomes to form large vesicles with a diameter up to 2 microns, named type I cytoplasmic vacuoles (CPV-I) where RNA synthesis continues ^43,52^. These modified endosomes formed after 3-6h of infection also support the translation of viral proteins and nucleocapsids ^52,54^. Here, we investigated the outcome of CHIKV ROs after 17 h of infection. This strategy revealed that CHIKV ROs are mainly located at the PM of HEK293T cells and attesting of the poor internalization capacity of these compartments as previously reported by others ^43^. We detected CHIKV ROs at the surface of the cell body and, also contrasting with SINV (José et al. 2017), on filopodia-like membrane protrusions were they colocalize with budding viral particles and with patches of membrane-anchored densities attributed to CHIKV envelope glycoproteins. This colocalization indicates the absence of exclusion process between spherules formation and particles budding sites. Interestingly, additional densities were found inserted in spherules delimiting membranes. While unexpected, recent immunolabelling studies revealed the presence of anti-E2 antibody staining at the surface of CHIKV spherules ^46^. Together with this observation, our results raise the question of the possible functional consequences of such incorporation. A direct cell-to-cell transport of budded particles to uninfected mosquito cells, involving the E2 envelope glycoprotein was previously reported ^55^. Whether it is tempting to speculate it could be a direct way for cell-to-cell transmission of CHIKV genetic material through filopodia, a recent hypothesis on the incomplete maturation state of these glycoproteins rather argues for a side effect resulting from the co-localization of replication and budding events, with no associated functional role. This question is therefore still open.

Surprisingly, CHIKV spherules imaged in the current study, whatever their localization on the PM, are heterogeneous in size, with a mean diameter of 90 nm, therefore contrasting with previous observations mainly performed from other model *alphaviruses* (i.e. SFV and SINV) or even for CHIKV for which ROs diameter was reported ranging from 50 to 70 nm at 6-8 hpi ^48^, a time corresponding to the end of the first viral cycle. Analyzing more closely CHIKV ROs size distribution revealed at least three main categories. ROs with maximal diameter of 84-100 nm represent >60% of total events, while compartments with a size of 40-83 nm or 101-152 nm were more rarely detected. As the size of ROs was correlated with RNA genome length contained in these compartments ^3,50^, and that the spherules of 50-70 nm were proposed to contained a single copy of the viral genome in double-stranded form ^48^, it signifies that the vast majority of CHIKV ROs observed at the PM at 17 hpi, corresponds to ROs maintained at the PM after the first viral cycle, continuing to replicate and accumulate viral RNA in their lumen. The variation of ROs diameters at the PM contrasts with homogeneity of spherule diameters in CPVs (50-60 nm). As the size of ROs in CPVs is very close to that of CHIKV ROs detected by others at the PM after 6-8 hpi ^48,49^, this argues for a regulated internalization mechanism, appearing at the end of the replication cycle, it means in the 6-8h after CHIKV infection. Consequently, we suppose that during the first 6-8h of infection, the spherules replicate RNA, growing up to the optimal size of about 50-70 nm. Then, a small part of these spherules is internalized in CPV, supporting the translation of viral proteins and nucleocapsids, likely explaining the presence of honeycomb arrangement capsid protein structures at vicinity of CPV (Figure S5). Therefore, the ROs internalization likely represents one pathway leading to the production of new infectious particles. However, it has also been shown that CHIKV can still replicate even if internalization of ROs is blocked ^56^. Consequently, this raises the question of the fate and possible role of CHIKV spherules staying at the PM after the first viral cycle.

As previously mentioned, the ROs observed at the PM at 17 hpi, corresponds to ROs maintained at the PM after the first viral cycle, whose RO diameters distribution superior to 80 nm reflects the continuous transcription of the various CHIKV RNA species in these compartments (i.e. (-)RNA, (+)RNA and subgenomic RNA). Determining the RNA species and copy number packaged in these various spherules remains a challenging issue to solve. Since SFV, SINV and CHIKV genomes are comparable in size, enlarged CHIKV ROs may reflect subtle differences in replication mechanisms. In this context, it would be also very interesting to explore the size of SFV or SINV ROs at longer infection times.

The presence of filamentous densities resembling RNA or ribosome-decorated RNA only extruded from ROs was observed only for ROs with diameters larger than 90 nm. Hence, this observation reinforces the idea of sequential models in which spherules would reach an optimal size, and therefore complete (-)RNA synthesis, before the replication complex switches its activity from (-)RNA synthesis to transcription and export of (+) stranded genomic and subgenomic RNAs ^25,43^.

Now, focusing on the architecture of ROs remained at the PM after the first viral cycle, cryo-EM and cryo-ET both revealed a density that separating the RO lumen from the cell cytoplasmic compartment. The sub-tomogram approach allowed us to compute a 3D reconstruction of the spherule neck, which forms a crown, a structural feature also found for FHV ^3,4^ or SARS-Cov-2 ^9^ replication organelles. It is worth noting that the fully symmetrical volume of the crown is not compatible with the fit of nsP1 and its partners (nsP2, nsp3 and nsP4) according to a 1:1 molecular ratio (nsP1: partner). This was confirmed by the cryo-EM single particle analysis of complexes assembled using recombinant CHIKV nsP1, nsP2 and nsP4 encoded by the related O’nyong-nyong *Alphavirus*, which assigned the central disk at the base of the neck crossed by an elongated density to the dodecameric nsP1 in interaction with the viral polymerase nsP4 facing the RO lumen and the nsP2 helicase-protease facing the cell cytoplasm ^49^ (Figure S7). The second ring of densities is supposed to correspond to nsP3 and associated cellular factors, including G3BP.

Thus, the structural organization of RO neck revealed in our study is in agreement with that described recently from infected cells after 6-8 hpi ^48,49^, meaning that the crown-like organization of the replication complex persists in ROs maintained at the PM even after 17 h of infection, arguing again for the functionality of ROs in our experimental conditions. Interestingly, the close inspection of ROs revealed that the crown organization of the complex is often difficult to see in individual sub-tomograms. Indeed, several patterns are mainly observed where all densities present in the tomographic reconstruction are not simultaneously present. Hence, the sub-tomogram averaging method provides an overview of the CHIKV replication complex at the base of the neck of spherules that is not completely realistic if the dynamical of the complex is not considered.

In conclusion of this study, we show that late CHIKV ROs are maintained at the PM beyond the first viral cycle, exhibiting a normally folded replication complex at the base of the neck, displaying an unexpectedly range of large RO diameters that reflects the role of these organelles in the viral RNA replication, with the ability also to export the viral RNA to the cytoplasm of cells where it is seemingly translated. All these points indicate that the late ROs are still active. This picture was not detected for ROs with a size below 50-70 nm. The observation of extruded RNA filamentous densities exclusively beneath ROs with a diameter greater than 90 nm supposes a sequential model in which spherules would reach an optimal final size before the replication complex switches its activity to (+) stranded genomic and subgenomic RNA transcription and export. In this dynamic picture, at the end of the first viral cycle, a few spherules of about 50-70 nm in diameter could be marginally internalized in CPVs according to a regulated process. At this time, spherules activity is switched from RNA transcription to its export and translation, resulting in the viral protein synthesis revealed by the presence of honeycomb arrangement of capsid proteins at vicinity of CPVs. Meanwhile, ROs maintained at the PM are actively involved in the production of new RNA genomes and new viral proteins required for assembly of nascent infectious viral particles, then also representing an alternative pathway if RO internalization failed. All these observations bring new insight into the CHIKV life cycle in human cells, highlighting the functional role of late ROs.

## EXPERIMENTAL PROCEDURES

### Materials

#### Cells

Human Embryonic kidney (HEK293T, ATCC #ACS-4500) and BHK-21 cells (ATCC #CCL-10) were used for viral propagation. African green monkey (Vero ATCC #CCL-81) cells were used for viral titration. All cells were cultured in Dulbecco modified Eagle medium (DMEM, Thermo Fisher Scientific) supplemented with penicillin, 10% foetal calf serum (FCS, Lonza) and grown at 37°C, in a 5% CO2 atmosphere.

#### Viruses

The pCHIKV-LR-5’GFP (LR-OPY1 isolate) and the CHIKV-377-mCherry (BNI-CHIKV_899 isolate) full-length molecular clones containing GFP and m-cherry reporters respectively ^33,40,57^ were transcribed *in vitro* using the mMESSAGE mMACHINE kit (Invitrogen, AM1344) following manufacturer’s instructions. RNA (1μg) was then transfected with Lipofectamine 2000 (Thermo Fisher Scientific) into 105 HEK293T cells and incubated at 37°C 5% CO2 for 24h. Supernatant was harvested, and virus stock was amplified using BHK-21 cells. After 48 hours culture supernatantwas collected, filtered through 0.45μm, aliquoted and stored at −80°C. Viral stocks were titrated using Vero cell plaque assays as previously described (Bernard et al., 2010). The virus assays were performed under BSL-3 safety conditions.

#### Immunofluorescence assay

CHIKV infected HEK293T cells grown on a glass coverslip were fixed with 4% formaldehyde/PBS (Sigma-Aldrich), permeabilized with 0.1% Triton X-100 in PBS, and blocked in PBS–2% FCS. Incubation with primary antibody was performed at 37°C for 1h at room temperature, and secondary reagents were added for 30 min at 37°C. Nuclei were stained with DAPI (4’,6’-diamidino-2-phenylindole; Sigma-Aldrich). After the final washes, coverslips were mounted with ProLong Gold antifade reagent (Invitrogen). Images were acquired using an Ayriscan superresolution microscope (Confocal Zeiss LSM880 Airyscan) at the Montpellier Resources Imaging (MRI) platform. Image processing and colocalization analysis were performed using ImageJ software ^58^.

#### Sample preparation - cryo EM

Cells were cultured directly on gold grids (R2/1, carbon film, Quantifoil) after being glow-discharged using a Pelco GD system. HEK293T cells were infected at the MOI of 50 for 17h in the growing medium and then 4% PFA fixed. After a brief wash with PBS, grids were loaded on the Mark IV Vitrobot system tweezers (FEI). Before 22 seconds of Whatman paper blotting at a −5-force offset, 3 μL of 10 nm BSA-treated fiducial gold particles (BBI solutions) were added on both sides of the grid. The chamber was kept under 100% humidity and at room temperature conditions. Grids were then rapidly plunged frozen in nitrogen-cooled liquid ethane and stored in liquid nitrogen waiting to be imaged.

#### Electron microscope setup – tilt series acquisition

For cryo-data acquisition, we used a Titan Krios cryo-TEM equipped with a field emission gun, operating at 300 kV (IRBA, Institut de Recherche Biomédicale des Armées, Brétigny-sur-Orge, France). Images were recorded on a Falcon III direct electron detector (Thermo Fisher Scientific). Regions of interest were recorded using MAPS software (Thermo Fisher Scientific). Tomographic tilt series were acquired with Tomo4 (Thermo Fisher Scientific) at a magnification of 29,000 × and exposed for 1.3-1.55 seconds, with a physical pixel size of 0.2931 Å and continuous focusing and tracking controls. Tomograms were collected with 2° tilt increment for a range spanning from −60° to +60° according to a dose symmetric acquisition tilt scheme ^59^, using defocus values ranging from −3 to −12 μm. The total electron dose per tomogram was between 120 and 140 electrons/Å^2^.

#### Tomographic reconstruction

In general, tomograms were reconstructed similarly as described elsewhere ^60^. Movies were firstly averaged and corrected using the *MotionCorr* software ^61^ . Tomogram reconstruction was then performed with Etomo from IMOD ^44^. The tilt series were coarsely aligned using cross correlation. CTF curves were estimated with *CtfPlotter* implemented in IMOD ^62,63^. To remove noise coming from high frequencies due to beam damages during tilt series acquisition, dose filtration was performed ^64^ and antialiasing filter was used. To this end, the *MtfFilter* function of IMOD was used, considering the cumulative dose of each image taken to apply increasing lowpass filters to consecutive images. The tomograms were reconstructed alternatively by weighted back-projection (WBP) and by SIRT (simultaneous iterations reconstruction technique) using 8 iterations and subsequently binned by 2, resulting in pixel sizes of 0.2931 nm (un-binned). To further facilitate visualization, a non-linear anisotropic diffusion (NAD) filter provided by IMOD is applied to the 2 times binned tomograms. Segmentation and animation were done using Amira software (Thermo Fisher Scientific).

#### Sub-Tomogram averaging

All sub-tomogram averaging steps were performed using the EMAN2 package ^65,66^ in version 2.9.9 unless stated otherwise. Masks were created with the EMAN2 e2filtertool.py and e2proc3d.py programs.

##### Pre-processing

Movie fractions were motion-corrected using MotionCor2 to compensate for beam-induced sample motion. A custom python script was used to automate that process and build a motion-corrected tilt-series stack, relying on the newstack and clip programs from the IMOD package ^44^.

##### Tilt-series alignment

A total of 38 tilt-series were aligned using EMAN2 landmark-based iterative approach and CTF estimation was performed. Low-pass filtered, binned by 4 tomograms were reconstructed for visualization and particle picking only.

##### Particle picking and extraction

We repurposed the e2tomo_drawcurve.py boxer program, originaly designed to pick particles along filaments to pick orientation aware spherule particles on bin 4 tomograms. Two points (3D coordinates) were assigned to each particle deemed suitable for analysis. One point was placed at the estimated center of the spherule neck and designates the center of the particle to be extracted. A second point was set inside the spherule compartment, towards its main axis as a way to manually assign an approximate orientation to the particle. Those coordinates were stored in json files. Stacks of per-particle CTF corrected 2D sub-tilt-series were extracted and corresponding 3D sub-tomograms reconstructed at binning 4 using e2spt_extract.py. The–curves option was set so as to read the particles orientation metadata and encode it in the extracted particle stacks header. The e2spt_extract.py script was modified so that it accepts particles with only 2 points defined and avoids doing any coordinates interpolation. A total of 463 binning 4 particles, 384 unbinned pixels in size were extracted from 38 tilt-series.

##### Initial model generation

A low-resolution initial model was generated at binning 8 using the iterative stochastic gradient descent (SGD) approach implemented in e2spt_sgd.py. Initial particle orientations read from their header were used as a seed for refinement. C57 symmetry was used during alignments and averaging as a way to improve the signal to noise ratio and force any rotational symmetry around the spherule main axis without applying any specific symmetry order. C1 refinements did not reveal obvious symmetry in the data, therefore we carried on using C57 symmetry for the downstream processing steps.

##### Model refinement and classification

Gold-standard single model refinements were done using the e2spt_refine_new.py program while multi-reference refinements employed e2spt_refinemulti_new.py. A first refined model was generated at binning 4 taking the initial model as a reference and using a large soft spherical mask, 192 unbinned pixels in diameter, for alignment. Particles orientations were refined within a search range of 16 degrees around their initial orientations determined at picking time. A first round of classification by iterative multi-reference refinement was performed using the same mask. Three references were initialized by randomly splitting particles in three sub-sets and averaging them. One class containing 166 particles was selected for further refinement. Particles belonging to that class were re-extracted at binning 2, with a size of 256 unbinned pixels, and a new model was refined from them. To tackle remaining heterogeneity in the data, a second round of classification without alignment was performed inside a tighter soft threshold based mask. Out of the four classes, the major class containing 98 particles was further refined inside a similar tight mask. For that last round of iterative refinement, both traditional iterations of 3D particles alignment refinement and 2D sub-tilt refinement including translation and rotation were performed.

##### Validation and post-processing

The gold-standard Fourier shell correlation (FSC) curve calculated from two independently refined half-maps using the same mask as the one used for refinement indicates an estimated resolution of 25 Å at FSC=0.143. The final map was Wiener filtered by the masked FSC.

#### High-Pressure Freezing and vitreous sectioning

The CHIKV-infected HEK293T cells were inactivated in 2,5% glutaraldehyde and 20% dextran-sodium cacodylate 0,1M. Pellets were frozen on copper tube carriers using the EM ICE high-pressure freezer (HPF, Leica). After HPF, the sample was processed by CEMOVIS (Cryo-Electron Microscopy of Vitreous Sections). Vitreous sectioning was performed on infected cells tubes containing the vitrified sample pre-cooled at −140°C on an EM UC6/ FC7 cryo-ultramicrotome (Leica Microsystems, Vienna). The sample was trimmed to a pyramidal shaped block of 140 μm base and approximately 50 μm height using a 45° cryo trimming diamond knife (CT441; Diatome, Biel, Switzerland). A cryo immuno diamond knife (MT15692; Diatome, Biel, Switzerland) with a clearance angle of 6° was used to get ribbons of cryo-sections at a nominal cutting feed of 50 nm and at cutting speeds of 40 mm/s. Ribbons of cryo-sections were attached to pre-cooled copper grids (hexagonal, 100 mesh with carbon and formwar), ready to be analysed under the Titan Krios.

## Acknowledgements

The work was supported by grant from the French Agence Nationale de la Recherche (ANR-18-CE11-002 to LB and PB). JG has a doctoral fellowship from ANR-18-CE11-002 and *la Fondation pour la Recherche Médicale* (FRM FDT202106013092). CBS is a member of the French Infrastructure for Integrated Structural Biology (FRISBI) supported by Agence Nationale de la Recherche (ANR-10-INBS-05).

